# Flexible backbone assembly and refinement of symmetrical homomeric complexes

**DOI:** 10.1101/409730

**Authors:** Shourya S. Roy Burman, Remy A. Yovanno, Jeffrey J. Gray

## Abstract

Symmetrical homomeric proteins are ubiquitous in every domain of life, and information about their structure is essential to decipher function. The size of these complexes often makes them intractable to high-resolution structure determination experiments. Computational docking algorithms offer a promising alternative for modeling large complexes with arbitrary symmetry. Accuracy of existing algorithms, however, is limited by backbone inaccuracies when using homology-modeled monomers. Here, we present Rosetta SymDock2 with a broad search of symmetrical conformational space using a six-dimensional coarse-grained score function followed by an all-atom flexible-backbone refinement, which we demonstrate to be essential for physically-realistic modeling of tightly packed complexes. In global docking of a benchmark set of complexes of different point symmetries — staring from homology-modeled monomers — we successfully dock (defined as predicting three near-native structures in the five top-scoring models) 19 out of 31 cyclic complexes and 5 out of 12 dihedral complexes.

**Highlights:**

- SymDock2 is an algorithm to assemble symmetric protein structures from monomers
- Coarse-grained score function discriminates near-native conformations
- Flexible backbone refinement is necessary to create realistic all-atom models
- Results improve six-fold and outperform other symmetric docking algorithms

**Graphical Abstract:** 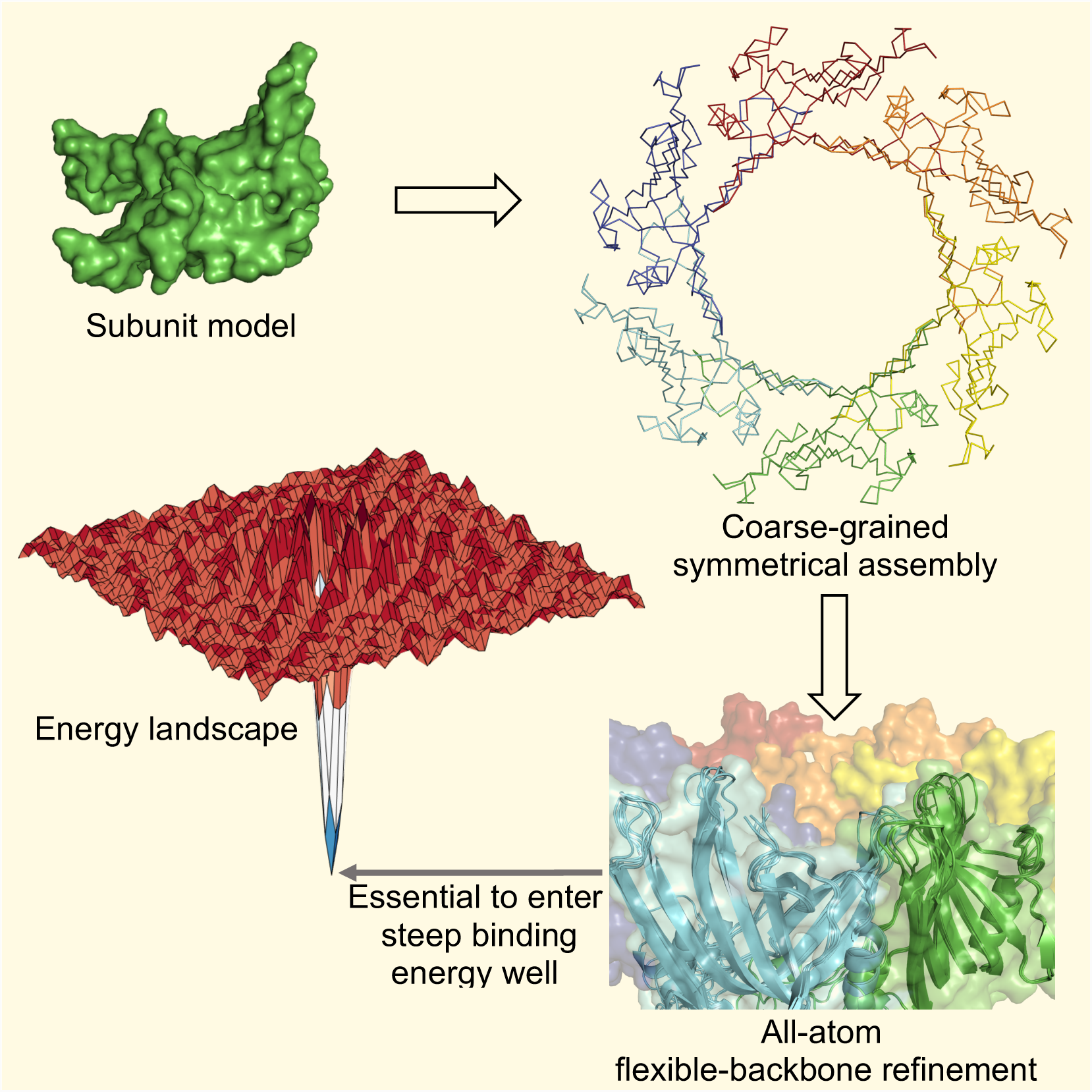

## Introduction

The pervasive appearance of symmetrical homomeric proteins across all domains of life has been attributed to increased stability (Wolynes, 1996), fine-tuned functional regulation (Monod et al., 1965; Perutz, 1989), better synthesis error control (Goodsell and Olson, 2000), reduced aggregation (Garcia-Seisdedos et al., 2017), and genome compactness (Crick and Watson, 1957). While these evolutionary forces drive proteins towards larger assemblies, the size of these complexes makes it particularly difficult to obtain high-resolution structures. Despite an estimated 50-70% of proteins being symmetric homomers (Levy et al., 2006), less than 40% of proteins in the Protein Data Bank (Berman et al., 2000) are symmetric (as of June 2018). This gap could be bridged by the development of computational docking methods to obtain accurate models of symmetric homomeric complexes.

Especially desirable are versatile methods that incorporate different kinds of experimental data in the modeling pipeline. For low-resolution cryo-EM, NMR or SAXS data, symmetry-constrained flexible refinement is essential for obtaining high-quality models (Chan et al., 2011; Joseph et al., 2016). In the absence of such a model, the method should be able to dock a homology modeled monomer (Eswar et al., 2006; Song et al., 2013; Yang and Zhang, 2015). This monomer model can be combined with experimental determination of the oligomeric state and/or symmetrical placement of subunits in homologs to prepare a preliminary model for refinement. If the relative orientations of the subunits cannot be obtained from homologous structures, the method should be able find the correct arrangement of the subunits while restricting the search space to relevant symmetrical conformations.

The symmetry framework in the Rosetta Macromolecular Modeling Suite allows modeling of complexes with arbitrary symmetries (DiMaio et al., 2011). The framework has been used to develop SymDock, a docking protocol for point symmetries. SymDock has been shown to correctly model complex structures from a monomer for a variety of symmetry groups (André et al., 2007). This protocol uses a coarse-grained phase to sample symmetric conformation space starting from a random or a pre-defined orientation followed by an all-atom phase for refinement. To further improve models, the Rosetta suite also allows integration of information from a plethora of experimental methods like cross-linking studies (Kahraman et al., 2013), NMR (Shen et al., 2008), and SAXS (Sønderby et al., 2017) as well as co-evolutionary analysis (Ovchinnikov et al., 2014) while docking. This two-phase approach and the variety of ways of adding constraints make SymDock an extremely versatile tool.

In the last published rounds of the blind docking challenge, Critical Assessment of PRediction of Interactions (CAPRI), although multiple groups generated high-quality models for various homodimers, no group was able to predict high-quality models for the five homotetramer targets, including two for which no acceptable solutions were submitted (Lensink et al., 2016). Recently, four leading symmetric docking methods were evaluated on a benchmark of 251 complexes, 180 of which were homodimers (Yan et al., 2018). Despite a favorable benchmark composition, starting with homology-modeled monomers, none of the methods was able to produce a CAPRI-acceptable model in the top ten predictions for more than half the complexes.

For Rosetta SymDock, we have been aware of two limitations preventing consistently accurate predictions. First, the scoring method used in the coarse-grained phase does not sufficiently discriminate between near-native and spurious interfaces. Second, in tightly packed complexes, small steric clashes between the subunits were not being resolved. Specifically, we observed that symmetric side chain packing and minimization were inadequate for resolving clashes between subunits in tightly packed complexes; such cases additionally require small backbone motions.

In this study, we address these limitations and demonstrate a drastically improved performance of this protocol. First, to enhance model evaluation in the coarse-grained phase, we employ a fast and accurate scoring scheme called Motif Dock Score (MDS). We previously developed MDS for docking heterodimeric complexes, and it greatly increased the number of conformations with near-native interfaces after a coarse-grained search (Marze et al., 2018). Second, we test two approaches to backbone flexibility that have been successfully used for heterodimeric complexes, *viz.* imitating conformational selection (Chaudhury and Gray, 2008; Moal and Bates, 2010; Venkatraman and Ritchie, 2012) and induced fit (Mashiach et al., 2010; Oliwa and Shen, 2015; Schindler et al., 2015). For conformational selection, we pre-generate an ensemble of conformations from the monomer and used them as input monomers for docking. For induced fit, we minimize energy along the backbone dihedrals and repack side chains during refinement, starting with a low repulsion between the atoms and progressively ramping it up. The refinement is performed after the rough subunit arrangement had been predicted in the coarse-grained phase.

We evaluate the enhanced protocol, SymDock2, on a diverse benchmark of 43 complexes belonging to the two most common symmetry groups, cyclic (described by a single rotational symmetry axis) and dihedral (described by a rotational symmetry axis and a perpendicular axis of two-fold symmetry). As these proteins rarely crystallize as monomers, we use monomers predicted by a homology docking server as a proxy for the ‘unbound’ structure. Given a particular point symmetry, we perform a global search of the relevant symmetrical conformation space. These inputs represent the most difficult case described earlier where the monomer conformation is approximate and the subunit arrangement is unknown. This workflow is similar to one commonly employed in CAPRI blind docking (Marze et al., 2017). The performance for both the coarse-grained and the all-atom phases show marked improvements over the original SymDock protocol without compromising the overall speed of the protocol.

## Results

Rosetta SymDock is a Monte Carlo-plus-minimization protocol (Li and Scheraga, 1987) that models symmetric homomeric complexes starting from a monomer structure and a symmetry definition (André et al., 2007). Symmetry definitions contain information about the rigid-body arrangement of the subunits, how to yield the energy of the whole complex from calculations on one subunit (or a set of subunits), and what the degrees of freedom are along which the subunits are allowed to move (DiMaio et al., 2011). For local docking, specific symmetry definitions can be recapitulated from a PDB file of a complex whereas for global docking, general symmetry definitions can be loaded for any given point symmetry. In the first, coarse-grained phase of the SymDock protocol, side chains are approximated as ‘pseudoatoms’. Coarse-graining allows the subunits to sample the symmetrical rigid-body conformations in a smoothened energy landscape. Next, the side chains are reintroduced and the putative encounter complex is refined by symmetrical side-chain optimization at the interfaces with minimal rigid-body motion. The protocol is illustrated as a flowchart in Fig. S1.

### Motif Dock Score discriminates near-native interfaces

First, we sought to produce low-scoring, near-native conformations by the broad, coarse-grained search of the symmetrical conformation space. To recognize a near-native conformation, the various interfaces between the subunits must be scored accurately. An ideal coarse-grained score function would recover the broad features of the all-atom energy landscape while smoothing over the local ruggedness.

The performance of the previous SymDock algorithm of André *et al* is shown in Figs. 1A, B, D, and E, which compare the docking landscapes after the coarse-grained phase and the full protocol for two example proteins, *viz.* Rhamnulose-1-phosphate aldolase (PDBID: 2V9N, symmetry: C4) and snRNP Sm-like protein (1H64, C7). Each model is the end-state of a global docking simulation and is represented as a point in terms of its deviation from the native conformation (root-mean-square deviation of C_α_ atoms) and its energy predicted by the given score function. The Rosetta all-atom score function (Alford et al., 2017; Park et al., 2016) scores models close to the native conformation more favorably than non-native models (Figs. 1A and D). However, the energy ‘funnels’ are absent for SymDock’s coarse-grained centroid score function (Figs. 1B and E, grey circles). The centroid score function does not score models under 5 Å RMSD_Cα_ any better than those far away from the native. Thus, the lowest-scoring structures in the coarse-grained phase are not useful input models for high-resolution refinement.

**Figure 1.**
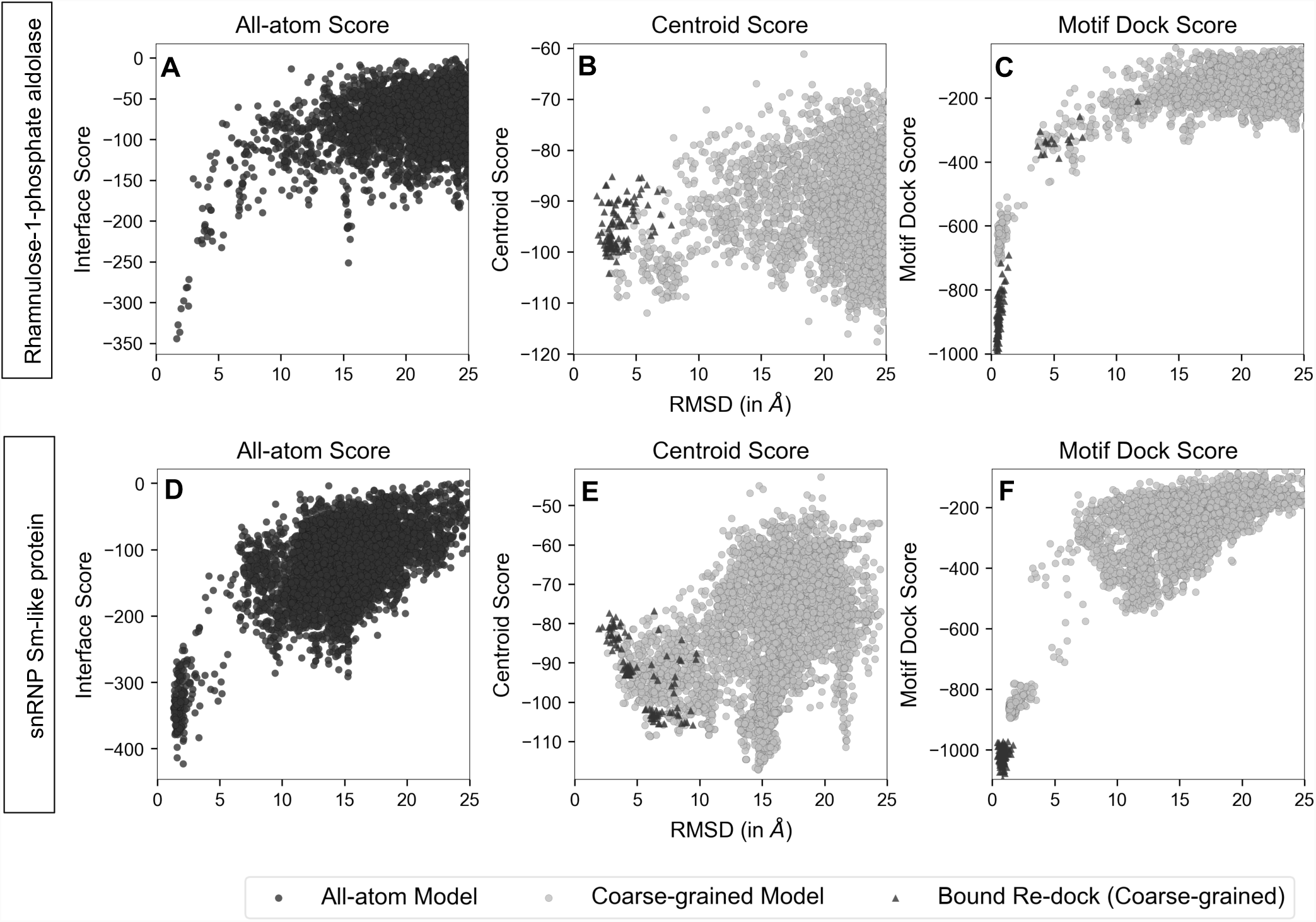
Comparison of energy landscapes in all-atom phase and coarse-grained phase. Score versus RMSD_Cα_ plots for two representative complexes, Rhamnulose-1-phosphate aldolase (A, B and C) and snRNP Sm-like protein (C, D and E) for 5,000 models generated from global docking of homology-modeled monomers (black, grey circles) and 100 models generated by redocking of bound subunits (triangles). Coarse-grained energy landscapes with Motif Dock Score (MDS) (C and F) resemble the all-atom energy landscapes (A and D), but those with Centroid Score (B and E) do not. Starting from the homology-modeled monomers, none of the 50 top-scoring models generated using Centroid Score are within 5 Å RMSD_Cα_. All the 100 top-scoring models generated using MDS are under 3 Å RMSD_Cα_. When re-docking bound subunits, closest models generated using Centroid Score (B and E) have 1.9 Å RMSD_Cα_and high relative scores in both cases. Bound re-docking with MDS (C and F) produces over 80% of the models docked to within 1 Å RMSD_Cα_ in both cases. These sub-angstrom re-docked models also score more favorably than all docking models made using homology-modeled monomers. Hence, Centroid Score does not recognize the energy well near the native conformation, whereas MDS does.

We considered the characteristics of the centroid score function to help identify opportunities to improve its accuracy. For the centroid score function environment and interacting residue pair terms, only the distance between backbone C_α_ atoms of two interacting residues across the interface is considered (Gray et al., 2003). Previous studies showed that this score function does not sufficiently discriminate near-native interfaces of heterodimers (Zhang et al., 2013). Also, to prevent favoring non-specific interactions across large, spurious interfaces, the residue-residue contact count of the centroid score function is capped, but this cap hinders the discrimination of large interfaces. Together, these score function features lead to the flat landscapes and false-positive energy wells observed in Figs. 1B and D.

Next, we tested whether better discrimination could be obtained by replacing the environment, pair, and contact scores with higher-resolution information about residue backbone orientation. For heterodimeric complexes, we recently developed Motif Dock Score (MDS), which radically improved coarse-grained interface detection (Marze et al., 2018). MDS is based on a residue-pair transform framework (Fallas et al., 2017). It estimates the minimum all-atom score for two residues interacting with a given backbone geometry defined by the six-dimensional transform (three rotations and three translations) required to superimpose the backbone atoms of one residue onto the other. To discretize the transform space, we use 2 Å grids for the translational dimensions and 22.5° grids for the rotational dimensions. For each residue type pair, we have pre-tabulated the lowest observed all-atom scores for every orientation present in high-resolution protein structures in the Protein Data Bank (Berman et al., 2000). If the orientation is not observed for that residue type pair (as is the case for the majority of the orientations), we score it as zero. To score a particular conformation in the coarse-grained phase, we look up the residue pair scores from these tables for every residue pair across the interface(s) of the principal subunit and sum them. The symmetry definition is then used to scale the score for the complex.

Figs. 1C and F (grey circles) shows that MDS scores near-native (<5 Å RMSD_Cα_) models better than far-away models. The general shape of the MDS energy landscape resembles that of the all-atom score function, with the aforementioned energy funnel near zero RMSD_Cα_. Moreover, of the 5,000 coarse-grained models obtained with MDS, 101 and 130 of them have RMSD_Cα_ values of less than 2 Å for Rhamnulose-1-phosphate aldolase and snRNP Sm-like protein, respectively, including 86 sub-angstrom models for the former. For comparison, there are no models within 2 Å RMSD_Cα_ for the coarse-grained phase with centroid score.

Another comparison of MDS score to the centroid score function is in the ranking of near-native models generated by re-docking the native assemblies. Ideally, the spread of RMSD_Cα_values should be minimal and the models should score better than the global docking models, as they represent the optimal solutions. Starting from the native configuration, centroid score forces the subunits to move away, with median RMSD_Cα_of 3.4 Å and 4.6 Å for Rhamnulose-1-phosphate aldolase and snRNP Sm-like protein, respectively (Figs. 1B and E, triangles). They also do not score better than the global docking models. In contrast, MDS scores them the lowest with median RMSD_Cα_ of 0.6 Å and 0.8 Å for the aforementioned complexes, respectively (Figs. 1C and F, triangles). Thus, MDS improves the docking performance of the coarse-grained phase, both in terms of the number of near-native models obtained and the ability to discriminate them.

Next, we expanded the comparison to a balanced benchmark of 43 complexes from the two most commonly found symmetry groups, cyclic and dihedral: five each for C2, C3, C4, C5, C6, D2, and D3 symmetries and two each for C7, C8, C9, and D4 symmetries. We challenged the methods with the hardest use-case, *viz.* no information is known apart from the sequence and the point symmetry. This test is akin to a round of the blind docking challenge, Critical Assessment of PRediction of Interactions (CAPRI), where no homologous complex exists for the modeling target. Starting from a homology-modeled monomer each complex, we generated 5,000 models (see Method Details and Table S1). In the 5, 50, and 500 top-scoring models, we counted the number of models within 5 Å RMSD_Cα_ of the native structure. Table 1 compares the bootstrapped averages for the coarse-grained phase run with centroid score and with MDS. For MDS, on average 1.96 of the 5 top-scoring models are near-native compared to 0.32 for centroid. MDS has a superior performance for the 50 and 500 top-scoring models as well.

**Table 1.**
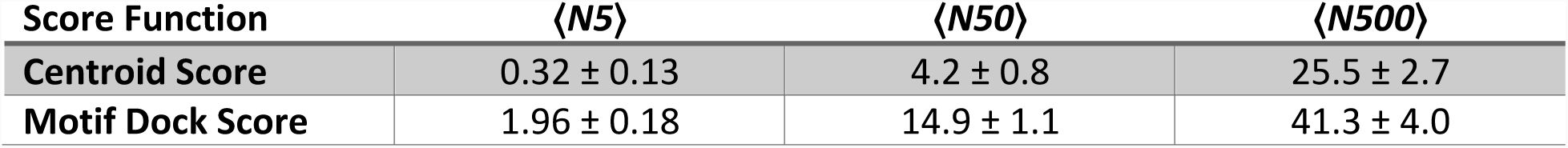
Average counts of near-native structures for the 5, 50, and 500 top-scoring models after the coarse-grained phase for coarse-grained score functions.

### Fixed-backbone refinement is insufficient to enter the binding funnel

After the initial arrangement of the subunits was calculated in the coarse-grained phase, we sought to produce a physically-realistic all-atom model. To do so, SymDock reintroduces the side chains, packs interface side chains, and refines the model by fixed-backbone energy minimization while allowing small rigid-body motions. With the coarse-grained phase using MDS producing accurate subunit arrangements, we presumed that this refinement would produce high-quality models. Surprisingly, models that were near-native after the coarse-grained phase had positive (unfavorable) interaction energy scores after refinement, indicating that the docked subunits scores worse than non-interacting monomers. Fig. 2A shows the MDS binding funnel for *Xenopus* Nucleophosmin (1XB9, C5) with models that have a positive post-refinement interaction score labelled red. About 22% of all structures are unfavorable, and all but one near-native (less than 5 Å RMSD_Cα_) are unfavorable. To confirm steric obstruction, we counted clashes as per the CAPRI definition (Méndez et al., 2003). Fig. S2 shows that in the 20 lowest-RMSD_Cα_models after fixed-backbone refinement, the average number of inter-chain atom-atom clashes is 50.6, compared to 21 in the native structure. We observed this insufficient refinement of near-native models for most complexes.

**Figure 2.**
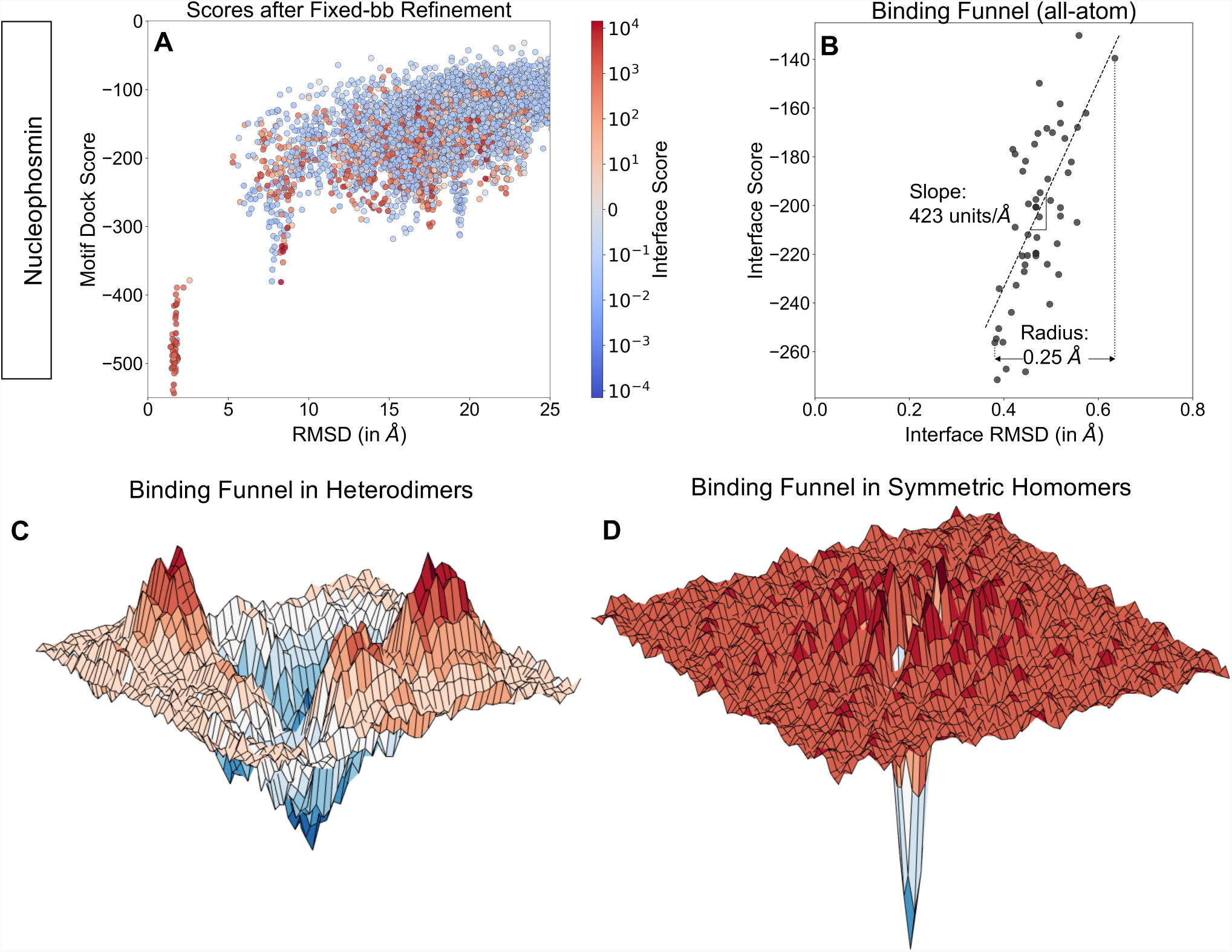
Fixed-backbone refinement is insufficient to enter narrow binding funnel. (A) Coarse-grained score versus RMSD_Cα_ (after coarse-grained phase) plots for *Xenopus* Nucleophosmin for 5,000 models. Models are colored by their interface score after fixed-backbone refinement. Almost all models under 5 Å RMSD_Cα_ have a positive interface after fixed-backbone refinement arising from minor clashes due to the introduction of side chains, despite repacking. Consequently, these models are discarded. (B) Interface score versus RMSD_Cα_ (after full-protocol) plots for *Xenopus* Nucleophosmin. A rapid drop in interface score between 0.6 and 0.4 Å RMSD_Cα_leads to an energy funnel with steep slope (dashed line) of 423 Å^−1^and a radius of 0.25 Å. (C) Conceptual representation of the energy landscape near the binding funnel for heterodimers. The funnel is comparatively shallow with local minima near it. (D) Conceptual representation of the energy landscape near the binding funnel for homodimers as seen by symmetrical docking protocols. The funnel is narrow and steep with no comparable local minima.

To test whether the refinement protocol works with an amenable backbone conformation, we generated 100 all-atom models starting from the native structure of Nucleophosmin. The average number of inter-chain clashes after refinement was 18.7, which was significantly lower than that of the global docked models (Fig. S4). All models were under 0.7 Å RMSD_Cα_ from the native structure and had highly favorable interaction energies (Fig. 2B). Since (a) the coarse-grained phase produces near-native subunit arrangements, and (b) fixed-backbone refinement can discover the binding funnel with the right backbone conformation, backbone errors in the homology-modeled monomers were likely causing the clashes in the docked models.

For the global docking simulations, four of the five homology-modeled monomers had backbones under 0.4 Å RMSD_Cα_, which was sufficient for assembling the subunits at the coarse-grained level, but insufficient for avoiding steric clashes with the side chains present. [In heterodimer docking, a monomer backbone with RMSD of 0.6 Å is typically sufficient for docking (Kuroda and Gray, 2016).] We speculated that when symmetry is enforced on an all-atom model, the leeway for backbone variation is markedly reduced. Minor deviations from the native backbone result in substantially higher energies as exemplified in Fig. 2B where a drop of 117 energy units takes place in 0.25 Å RMSD_Cα_

Compared to heterodimers where the average binding funnel slope is 15 Å^−1^the slope for this complex was unusually steep. For the homomeric complexes in our benchmark, we found the average slope of the binding funnel to be 249 Å^−1^. Further, the average radius of the binding funnel was found to be 0.26 Å for these complexes as opposed to 0.41 Å for heterodimers (see Quantification and Statistical Analysis). These observations are conceptually represented in Figs. 2C and D, where homomers have a narrower, steeper well in the rugged all-atom energy landscape as compared to heterodimers. More examples of binding funnel data for homomers and heterodimers can be found in Fig. S3. As homomers generally have extensive interfaces owing to multivalent interactions, we normalized the slopes by dividing them by the lowest interface score observed for the complex to obtain slopes of 0.62 Å^−1^. and 0.31 Å^−1^., respectively for homomeric and heterodimeric complexes. Even after normalization, funnels in homomers are twice as steep as heterodimers. We concluded that in homomers, for a backbone with errors, no amount of side chain packing can help it find the narrow binding funnel. Flexible-backbone strategies are required to reduce steric clashes and build physically-realistic models.

### In context, flexible backbone refinement is crucial to enter the binding funnel

To find alternative routes to enter the binding funnel, we tried mimicking natural mechanisms of backbone flexibility. Two kinetic mechanisms widely observed in assembly and regulation of proteins are conformational selection and induced fit (Changeux and Edelstein, 2011).

### Imitating Conformational Selection

We have previously leveraged conformational selection to improve the docking performance of heterodimeric complexes by pre-generating an ensemble of backbone conformations from the individual monomers and docking the optimal backbones (Marze et al., 2018). Using a similar approach, starting from a homology-modeled monomer, we generated 50 conformers each using three conformer generation methods: perturbations along the normal modes by 1 Å (Go et al., 1983), small backbone perturbations using Backrub (Smith and Kortemme, 2008), and general refinement using Rosetta’s Relax protocol (Tyka et al., 2011) (see Method Details). We supplemented the ensemble of the five original homology-modeled backbones with the new 150 backbone conformations. We ran 500 independent fixed-backbone simulations with each of the 155 monomer backbones and bootstrapped the results to simulate the selection of 2,500 models (see Method Details and Quantification and Statistical Analysis).

Next, we tested the efficacy of starting with these large, diverse ensembles using a small benchmark of 10 cyclic complexes. We compared the number of structures with RMSD_Cα_ less than 5 Å from the native in the 1% top-scoring models, i.e. the 25 top-scoring models. Fig. 3 shows a case-by-case comparison. Docking with just the homology models (HM/Fixed-bb) gives a median value of 9.6 near-native models after the coarse-grained phase, which goes down to 3.0 after the full protocol. Using a mixture of conformations (HM+Ens/Fixed-bb), the results get marginally worse with median values of 6.8 and 2.8 near-native models, respectively, for the coarse-grained phase and the full protocol. Starting with a large ensemble improves performance for some complexes and makes it worse for others. In general, backbone conformations generated from the monomer lack information about where the other subunits are and encounter the same barriers as the original homology models. Thus, our conformational selection approach was unable to improve docking accuracy for symmetric homomers.

**Figure 3.**
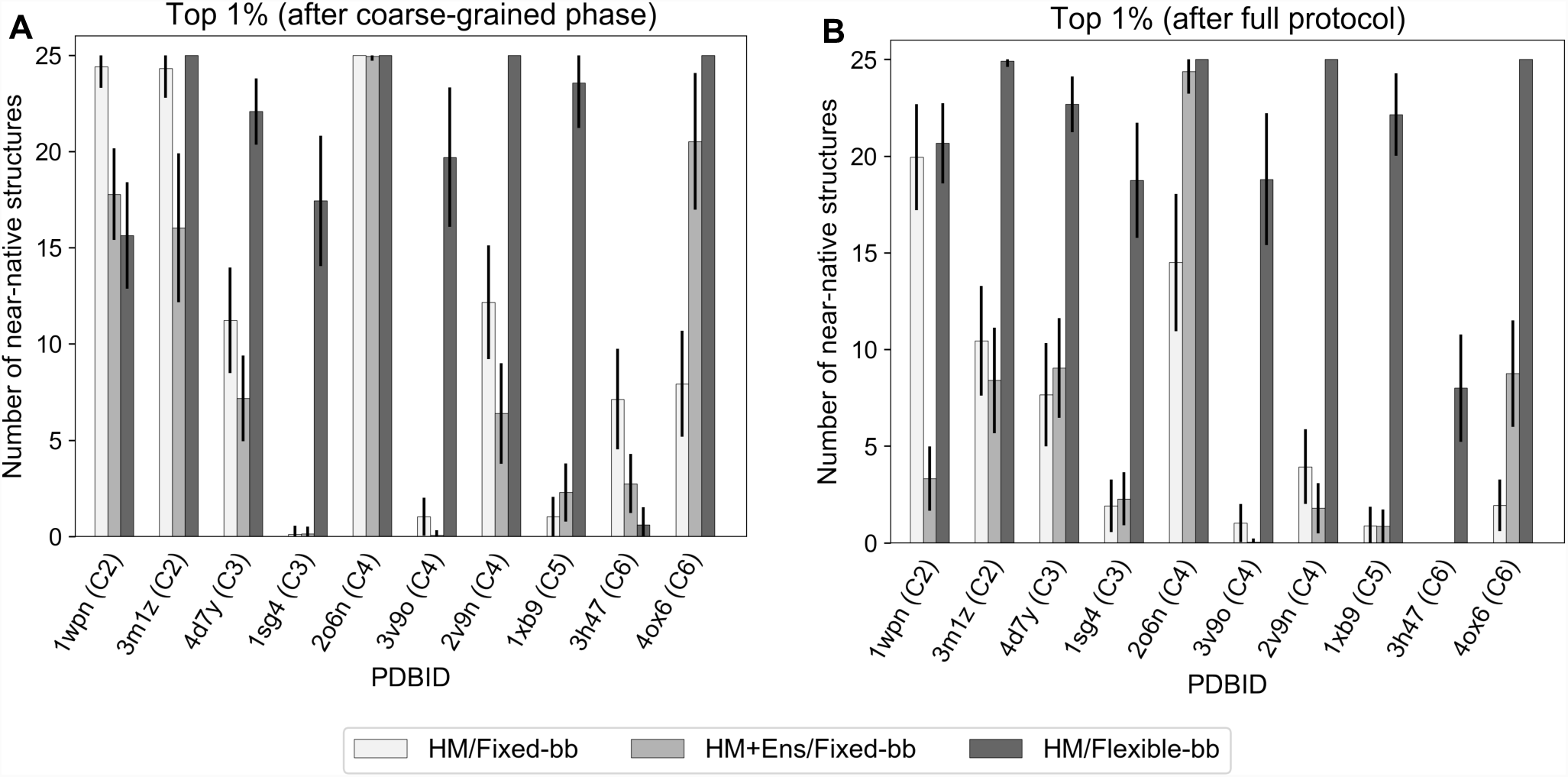
Flexible-backbone refinement improves docking performance. Comparison of bootstrapped averages of the number of near-native structures in the set of 2,500 docking models using: [white] the homology models (HM) and fixed-backbone refinement, [light grey] homology models supplemented with an ensemble of 150 pre-generated backbone conformations (HM+Ens) and [dark grey] fixed-backbone refinement, and the homology models and flexible-backbone refinement after the coarse-grained phase (A) and after the full protocol (B). Starting with 150 additional backbone conformations generated without the context of the complex improves docking performance for 4 out of 10 complexes, but makes it worse for 5 complexes. Starting with just the homology models and performing flexible-backbone refinement leads to improvements in 9 out of 10 complexes after the coarse-grained phase and in all complexes after the full protocol. After flexible-backbone refinement, more than 70% of the top-scoring models were near-native for 9 out of the 10 complexes.

### Imitating Induced Fit

We next hypothesized that the backbone needed to be adjusted in the context of the complex and not independent of it. That is, since the coarse-grained phase was correctly predicting the rigid-body arrangement of the subunits, we tested using these coordinates to induce a backbone fit at the interface. Specifically, we alternately repacked side chains and minimized the energy of the whole protein while slowly ramping up the repulsive component of the van der Waals potential from 2% to 100%. Gradient-based energy minimization along the backbone dihedral angles (ϕ and ψ) provided an avenue for the backbones to relieve clashes. The presence of the other subunits provided the necessary constraints to move the backbone to best fit the complex. To ensure a constant context, we removed rigid-body moves. Starting with just the five homology-modeled monomers per complex, we generated 5,000 docked models.

Finally, we tested this approach for the same benchmark of ten proteins (HM/Flexible-bb) and bootstrapped the results to simulate the selection of 2,500 models (see Quantification and Statistical Analysis). In the top-scoring 1% of models, the median counts of near-native models increased to 21.1 after the coarse-grained phase and 22.3 after the full protocol (Fig. 3). Thus, we conclude that inducing a change in the backbone retains good coarse-grained models and gains additional near-native models for all complexes tested. Further, the average number of inter-chain clashes in the 20 lowest-RMSD_Cα_models decreases from 50.6 in fixed-backbone refinement to 14.5 (Fig. S4).

### Improvement in global docking performance over a diverse benchmark

In the two-stage Rosetta SymDock2 protocol, we combine the coarse-grained phase with MDS with an in-context, flexible-backbone, all-atom refinement. To evaluate the performance of Rosetta SymDock2 and compare it to SymDock across a benchmark of 43 proteins, we performed a global docking search along symmetrical conformation space starting from five homology-modeled input monomers per target to generate 5,000 candidate models for each complex. Next, we resampled the docked models and reported averages and medians for targets success metrics based on the near-native model counts. For the coarse-grained phase, we defined a near-native model as one with RMSD_Cα_ under 5 Å. For the full protocol, we defined near-native as acceptable, medium-quality, or high-quality as per the CAPRI criteria, which are based on the ligand RMSD_bb_, interface RMSD_bb_, and fraction of native contacts recovered (Méndez et al., 2003) and detailed in Quantification and Statistical Analysis.

To test near-native sampling and discrimination ability, we counted the number of near-native models in the five top-scoring models and averaged over resampling attempts to calculate the ⟨*N5*⟩ metric (see Quantification and Statistical Analysis). ⟨*N5*⟩ after the coarse-grained phase indicates the ability of the broad search in the coarse-grained space to find approximate solutions. Most importantly, ⟨*N5*⟩ after the full protocol determines the overall accuracy of the method. For SymDock2, the average ⟨*N5*⟩ value improved from 2.0 to 2.8 going from the coarse-grained phase to the full protocol, indicating that while the broad search in the coarse-grained space found approximate solutions, the introduction of side chains and flexible-backbone refinement further discriminated near-native models. SymDock had an average ⟨*N5*⟩ value of 0.3 for both the coarse-grained phase and 0.8 for the full protocol suggesting a failure to sample and discriminate near-native models in most complexes. Fig. 4 presents a case-by-case comparison between the two methods, and Table 2 provides a category-wise summary of the benchmark results. The average performance on cyclic complexes is better than on dihedral complexes on every metric. Detailed metrics and plots for each complex can be found in Table S2 and Figs. S5– S8.

**Table 2.** Category-wise summary of the results of Rosetta SymDock and SymDock2 across a benchmark of 43 complexes.

|  | Category | SymDock |  |  | SymDock2 |  |  |
| --- | --- | --- | --- | --- | --- | --- | --- |
| | | Coarse-grained<br>$\langle E_{1\%} \rangle$ | Coarse-grained<br>$\langle N5 \rangle$ | Full protocol<br>$\langle N5 \rangle$ | Coarse-grained<br>$\langle E_{1\%} \rangle$ | Coarse-grained<br>$\langle N5 \rangle$ | Full protocol<br>$\langle N5 \rangle$ |
| Average Value <sup>a</sup> | **Cyclic** ($n = 31$) | 6.7 | 0.4 | 1.1 | 34.6 | 2.6 | 3.1 |
| | **Dihedral** ($n = 12$) | 6.4 | 0.1 | 0.2 | 15.8 | 0.2 | 1.8 |
| | **All** ($n = 43$) | 6.6 | 0.3 | 0.8 | 29.3 | 2.0 | 2.8 |
|  | **All w/ 1% filter** <sup>c</sup> | N/A | N/A | 1.0 | N/A | N/A | 2.9 |
| Expected Success <sup>b</sup> | **Cyclic** ($n = 31$) | 6 | 2 | 4 | 14 | 16 | 19 |
| | **Dihedral** ($n = 12$) | 0 | 0 | 0 | 0 | 0 | 5 |
| | **All** ($n = 43$) | 6 | 2 | 4 | 14 | 16 | 24 |
|  | **All w/ 1% filter** <sup>c</sup> | N/A | N/A | 4 | N/A | N/A | 25 |

<sup>a</sup> Values are the average of bootstrapped means across all complexes of the specified category.
<sup>b</sup> Expected success is the number of successfully-docked complexes based on the following criteria: for $N5$ , $\langle N5 \rangle \geq 3$ ; for $E_{1\%}$ , $\langle N50 \rangle \geq 15$ .
<sup>c</sup> Only the 1% top-scoring models after the coarse-grained phase underwent all-atom refinement.

**Figure 4.**
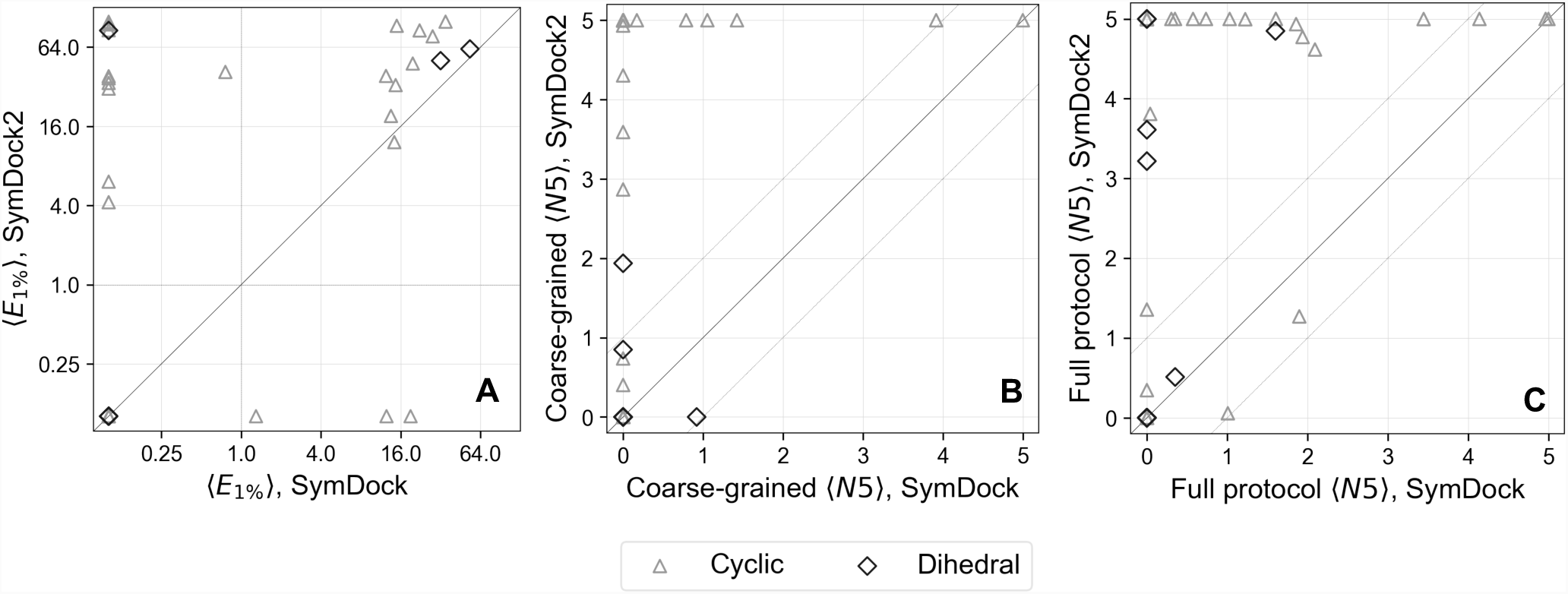
Rosetta SymDock2 compares favorably with SymDock on various assessment metrics. Comparison of bootstrapped-averaged metrics for 43 individual complexes (31 cyclic complexes [triangle] and 12 dihedral complexes [diamond]) both after the coarse-grained phase (A and B) and after the full protocol (C) shows significant performance gains. All complexes (points) above the diagonal line are improved in SymDock2. (A) Comparison of fold-enrichment of near-native models in the 1% top-scoring models, ⟨*E1%*⟩ on a log-log scale shows a higher enrichment in 19 cyclics and 3 dihedrals and a lower value in 4 cyclics and 0 dihedrals. Complexes to the right of the vertical dashed line are enriched in SymDock, and complexes above the horizontal dashed line are enriched in SymDock2. (B and C) Comparison of number of near-native models in the five top-scoring models, ⟨*N5*⟩, shows marked improvements both after the coarse-grained phase (B) and after the full protocol (C). Areas above and below the dashed lines indicate cases where the two methods differ significantly, *i.e.* by more than 1 model on average. SymDock2 has significant improvements in 16 cyclics and 1 dihedral complex after the coarse-grained phase, and most importantly, in 17 cyclics and 5 dihedrals after the full protocol. No complexes were modeled significantly worse with SymDock2.

We classified a homomeric complex as successfully docked if ⟨*N5*⟩ ≥ 3, i.e. at least 3 of the 5 top-scoring models are near-native on average. This criterion indicates that the protocol converges on the native structure. While SymDock docked 4 of the 43 complexes successfully, SymDock2 docked 24 of them successfully, representing a six-fold improvement in the success rate of blind docking for a general case. We observed performance gains for both symmetry groups, with 15 new cyclic complexes and 5 new dihedral complexes being docked successfully.

To estimate how many independent trajectories must be run to completion, we evaluated the fold-enrichment of near-native models for the top-scoring 1% of models after the coarse-grained phase, ⟨*E*_*1%*_⟩. SymDock2 had an average ⟨*E*_*1%*_⟩ value of 29.3, indicating a highly enriched low-scoring model set, while SymDock had a lower average ⟨*E*_*1%*_⟩ value of 6.6. Thus, if we were to only refine the top-scoring 1% of models after the coarse-grained phase of SymDock2, the average ⟨*N5*⟩ value after the full protocol would be 2.9. Furthermore, the number of complexes successfully docked for SymDock2 increases to 25. To explain the increase in success rates, we consider the example of Acylhomoserine lactonase (4ZO2, C2). With the all-atom score function, a non-native binding mode around 10 Å RMSD_Cα_ from the native is scored more favorably than near-native conformations (Fig. S7). MDS discriminates the native biding mode better and no models having the aforementioned non-native binding mode are selected in the top 1% (Fig. S5). By reducing the number of false positives, the success rates are increased. Thus, we recommend running SymDock2 as a two-step protocol where only the top-scoring 1% of coarse-grained models are refined (see flowchart in Fig. S2).

### Flexible-backbone refinement does not affect net efficiency

Compared to fixed-backbone refinement, modeling backbone motions requires sampling an exponentially larger conformational space. Instead of explicitly sampling backbone changes, we employed systematic energy minimization along backbone torsions to induce a fit. In fixed-backbone refinement of the interfaces, the computational time depends on the interface size and is largely independent of the monomer size. On the other hand, energy minimization along the backbone involves small changes in the subunit core to better accommodate the interfaces and hence, the time increases with monomer size. In fact, SymDock2 was between 2 and 3 times slower than SymDock for models that had larger interfaces to be fit (data not shown). Flexible backbone refinement also led to the fitting of spurious interfaces that were then weeded out by their relatively poor interface scores. As a result, almost every SymDock2 model had a negative interface score. Compared to SymDock, where a significant number of models are filtered out because of positive interface scores, induced fit reduces the total number of models rejected with the same filters (see Method Details for filter values). As a result of the low rejection rate, SymDock2 can compensate for the additional time required for flexible-backbone refinement by attempting fewer trajectories. We have previously shown that MDS is marginally faster than centroid score in the coarse-grained phase (Marze et al., 2018), which too works in favor of SymDock2. For an even comparison, in Fig. 5, we show the time per model for the two methods for every complex when all coarse-grained models are refined. SymDock was faster for 22 complexes and SymDock2 for 20 complexes. SymDock was typically faster for larger complexes and SymDock2 for smaller complexes. In 31 of the 43 complexes the run time difference was less than ± 20%, with the largest difference being less than 70%.

**Figure 5.**
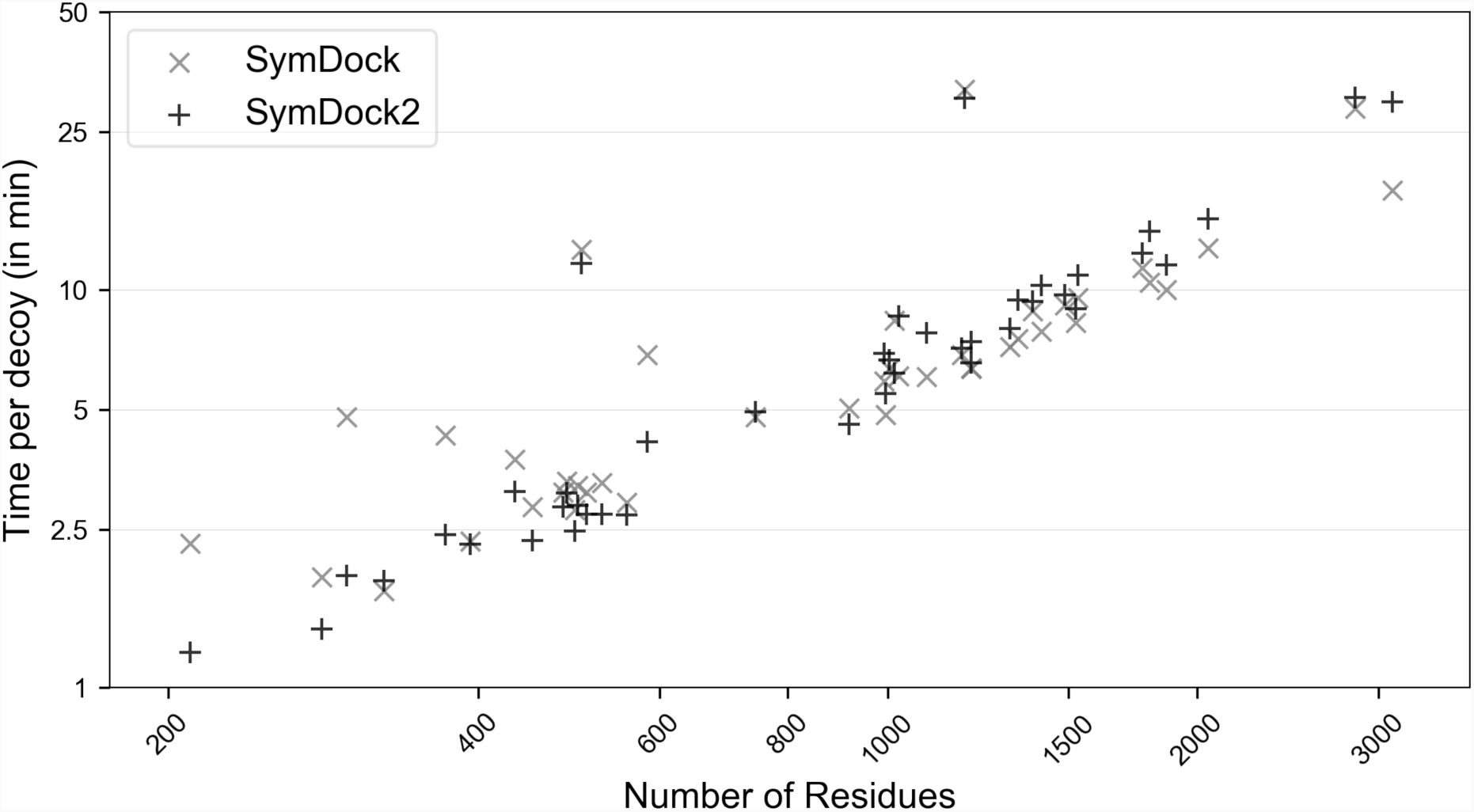
On average, Rosetta SymDock and SymDock2 have similar per-decoy runtimes in the benchmark. Comparison of average time per decoy on a log-log plot demonstrates similar scaling with complex size and symmetry for SymDock (×) and SymDock2 (+). Despite having a slower all-atom refinement phase, no complex had a more than a 70% overhead with SymDock2. For the two methods, run times were within ± 20% for 31 out of 43 complexes.

High fold-enrichment of near-native models for the 1% top-scoring models after the coarse-grained phase allowed us to considerably reduce the number of models refined using the expensive all-atom refinement. Broad sampling in the coarse-grained phase takes 51-78% of the time in each trajectory. By carrying forward only the top 1% from the coarse-grained phase to the refinement phase, we could save 22-49% of the total time. For the average complex in our benchmark, to generate 5,000 coarse-grained models and then refine 50 of them, SymDock2 requires 89 hours on a 4-core personal computer or under 1 hour on a 360-core cluster.

## Discussion

Here, we have developed and benchmarked a method to accurately model homomeric assemblies from an approximate monomer structure and the point symmetry. Our first innovation was using a six-dimensional coarse-grained scoring scheme, MDS, to successfully discriminate near-native interfaces with accuracy comparable to an all-atom score function. The second innovation was refining approximate models with small backbone motions to fit tight complexes together. Taken together, these two advances achieve successful blind global docking of six times as many complexes as Rosetta’s original SymDock (André et al., 2007). In Table 3, we compare the global docking performance of Rosetta SymDock and the new SymDock2 to four leading homomer docking methods recently tested by Yan *et al*: SymmDock (Schneidman-Duhovny et al., 2005), M-ZDOCK (Pierce et al., 2005), SAM (Ritchie and Grudinin, 2016), and HSYMDOCK (Yan et al., 2018). In this table, to compare to the other methods, we changed our success criterion to match their criterion of ⟨*N10*⟩ ≥ 1, *i.e.* at least one of the ten top-scoring models should be CAPRI acceptable, medium-or high-quality. While the methods are tested on different benchmarks with different ways of generating unbound structures, general patterns can be observed. With a success rate of 71% for cyclic complexes, Rosetta SymDock2 outperforms other methods. For dihedral complexes, SymDock2’s success rate of 50% is comparable to HSYMDOCK. Moreover, flexible refinement in SymDock2 ensures that the interfaces are relatively free of clashes, which were frequently observed with fixed-backbone docking of tightly-packed homomers.

**Table 3.** Comparison of leading symmetrical homomer docking methods with Rosetta SymDock2.

| Method | Description | Accuracy |  |
| --- | --- | --- | --- |
|  |  | Cyclic Complexes | Dihedral Complexes |
| SymmDock<br>(Schneidman-Duhovny et al., 2005)<sup>a</sup> | Local feature matching, cluster evaluation | 79/213<br>(37.1%) | N.A.<sup>d</sup> |
| M-ZDOCK<br>(Pierce et al., 2005)<sup>a</sup> | FFT docking, model evaluation | 86/213<br>(40.4%) | N.A.<sup>d</sup> |
| SAM<br>(Ritchie and Grudinin, 2016)<sup>a</sup> | Spherical polar FFT docking, model evaluation | 94/213<br>(44.1%) | 13/35<br>(37.1%) |
| HSYMDOCK<br>(Yan et al., 2018)<sup>a</sup> | FFT docking, cluster evaluation | 104/213<br>(48.8%) | 19/35<br>(54.3%) |
| Rosetta SymDock<br>(André et al., 2007)<sup>b,c</sup> | Monte Carlo docking, model evaluation | 15/31<br>(48.4%) | 2/12<br>(16.7%) |
| Rosetta SymDock2<br>(This article)<sup>b,c</sup> | Monte Carlo docking, model evaluation | 22/31<br>(71.0%) | 6/12<br>(50.0%) |

<sup>a</sup> Benchmark set: Yan et al., 2018
<sup>b</sup> Benchmark set: this article
<sup>c</sup> For an even comparison of all the methods, the metric for success was changed from $\langle N5 \rangle \geq 3$ to $\langle N10 \rangle \geq 1$ .
<sup>d</sup> Dihedral complex docking is not available with this method.

We also explored some characteristics of the interfaces of homomers and compared them to those of heterodimers. We developed MDS with the conjecture that given the relative orientation between backbones of interacting residues, we can estimate the optimal side chain interaction energy. The broad bin size of the score tables prevents overfitting to any particular protein class (Marze et al., 2018). Although MDS was developed to recognize hetero-oligomeric interfaces, it performs just as well for homomers, which suggests that at the level of individual residue pairs, the interfaces of homo-and hetero-mers have similar interactions.

We found that the near-native binding energy landscape was 14 times steeper on average than that of heterodimers. Even after normalizing for the depth of the binding funnels, the homomer funnels were twice as steep as those of heterodimers. This energy landscape is seen by symmetric docking protocols with enforced symmetry, which is not a constraint for the natural association of the subunits for symmetries higher than C2. For example, D2 complexes likely assemble as dimer of dimers with the ratio of the interaction strengths of the different interfaces dictating the hierarchy of assembly (Villar et al., 2009). For some proteins, inter-subunit interactions may be essential to find an energy funnel while folding (Wolynes, 1996), and hence an independent docking landscape may not exist. Reproducing these multi-state interactions becomes infeasible for a general case where the pathway of association is unknown, and so we resort to docking all the subunits together symmetrically. However, once assembled, the depth of the energy funnel suggests that symmetry confers stability through multivalent associations.

Most of the proteins that we have considered in the benchmark are globular, which allowed us to deconstruct the problem into generating an approximate monomer and then docking and refining it. The homology server we used, Robetta, performed admirably with monomer modeling. For the few cases for which it did not produce a monomer model under 2 Å RMSD, our docking performance suffered. For example, for phage SF6 terminase small subunit (3ZQO), the complex is stabilized by intertwined interfaces, and hence Robetta failed to create a good monomer model. Such proteins require simultaneous folding and docking. Previous fold-and-dock attempts have achieved success rates similar to that of symmetrically docking small globular proteins (Das et al., 2009). However, owing to the sheer size of the conformational space that needs to be sampled, without experimental constraints, *de novo* folding and docking is currently feasible only when subunits are smaller than 100 residues. Instead, incorporating symmetry information while homology modeling provides a promising avenue, which servers like Robetta (Park et al., 2018), SWISS-MODEL Oligo (Bertoni et al., 2017) and GalaxyHomomer (Baek et al., 2017) have recently demonstrated. Unfortunately, >90% sequence identity is required to guarantee symmetry type conservation (Levy et al., 2008); at <30% identity, interactions may differ completely (Aloy et al., 2003). In case of Robetta, 11 of the 22 complexes of the CASP12 experiment (Kryshtafovych et al., 2018) did not have sufficient symmetric templates and in case of SWISS-MODEL Oligo, 20% of the complexes considered did not have any viable symmetric templates. Thus, for large homomers, especially for those without close homologs, symmetric docking methods are required for modeling the complex.

The versatility of the techniques developed here facilitates application across a broad spectrum of problems. Integration with Rosetta’s input system allows us to incorporate cryo-EM, NMR, SAXS, cross-linking, and sequence co-evolution data (Kahraman et al., 2013; Ovchinnikov et al., 2014; Shen et al., 2008; Sønderby et al., 2017). In combination with these data, SymDock2 is a powerful tool for understanding homomer assembly and function.

## Author Contributions

Conceptualization and Methodology, S.S.R.B. and J.J.G.; Investigation, S.S.R.B. and R.A.Y.; Writing S.S.R.B., R.A.Y, and J.J.G.; Funding Acquisition, Resources, and Supervision, J.J.G.

## Acknowledgements

This work has been supported by grants from the National Institutes of Health, USA (grant R01-GM078221) and the National Science Foundation, USA (award 1507736). Computations in this study have been performed in part on the Maryland Advanced Research Computing Center (MARCC) Blue Crab cluster. The authors thank Prof. Ingemar André, Lund University for helpful discussions and guidance and Dr. David E. Kim, University of Washington for advice on homology modeling with Robetta.

## STAR٭Methods

### Key Resources Table

| Software and Algorithms |  |  |
| --- | --- | --- |
| Rosetta SymDock | André et al., 2007 | https://www.rosettacommons.org/software |
| Rosetta SymDock2 | This article | https://www.rosettacommons.org/software |
| Rosetta Relax | Tyka et al., 2011 | https://www.rosettacommons.org/software |
| Rosetta Backrub | Smith and Kortemme, 2008 | https://www.rosettacommons.org/software |
| RosettaScripts | Fleishman et al., 2011 | https://www.rosettacommons.org/software |
| PyRosetta | Chaudhury et al., 2010 | http://www.pyrosetta.org/ |

## Method Details

### Benchmark set generation

We generated a benchmark of symmetric homomeric proteins from structures deposited in the Protein Data Bank (PDB) having either cyclic (C2-C9) or dihedral (D2-D4) point symmetries. First, we filtered structures by resolution, retaining those with a resolution of 1.5 Å or better for C2-C4 and D2-D4, with a resolution of 2.0 Å or better for C5-C7, and with a resolution of 2.5 Å or better for C8-C9. We then randomly chose complexes from each symmetry group and retained those that passed the following selection criteria. We discarded entries with atoms having B factors greater than 40 or ligands with more than 5 atoms at the interfaces. Additionally, we only included entries for which the biologically relevant symmetry is confirmed in a publication. Next, we discarded any entry for which the earliest `REVDAT` record date was earlier than 2002. Within each symmetry group, we selected complexes to include a range of monomer sizes and diversity in secondary structural elements. We did not filter out complexes with intertwined interfaces. In total, we selected a benchmark of 43 complexes of different symmetries: five each for C2, C3, C4, C5, C6, D2, and D3 symmetries and two each for C7, C8, C9, and D4 symmetries. The PDBs chosen were:

| Symmetric Benchmark Set |  |  |  |  |  |
| --- | --- | --- | --- | --- | --- |
| *Target* | *Symmetry* | *Resolution (Å)* | *Type* | *Subunit size (residues)* | *Earliest REVDAT* |
| 3urr | C2 | 1.4 | transferase | 153 | 21-Dec-11 |
| 1wpn* | C2 | 1.3 | hydrolase | 188 | 23-Nov-04 |
| 2nlv | C2 | 1.3 | Xisl-like (unknown function) | 112 | 5-Dec-06 |
| 3m1z* | C2 | 1.42 | lyase | 228 | 16-Jun-10 |
| 4zo2 | C2 | 1.09 | hydrolase | 294 | 15-Jul-15 |
| 1sg4* | C3 | 1.3 | isomerase | 260 | 18-Jan-05 |
| 4co0 | C3 | 1.4 | signaling protein | 112 | 28-May-14 |
| 4d7y* | C3 | 1.44 | signaling protein | 146 | 28-Jan-15 |
| 4xla | C3 | 1.47 | viral protein | 600 | 20-Jan-16 |
| 5i6n | C3 | 1.22 | oxidoreductase | 332 | 13-Jul-16 |
| 1p5b | C4 | 1.35 | oxidoreductase | 380 | 28-Oct-03 |
| 2o6n* | C4 | 1.1 | de novo protein | 35 | 23-Oct-07 |
| 2v9n* | C4 | 1.4 | lyase | 274 | 15-Jan-08 |
| 4z0g | C4 | 1.25 | oxidoreductase | 413 | 25-Nov-15 |
| 3v9o* | C4 | 1.45 | lyase | 143 | 25-Jan-12 |
| 1xb9* | C5 | 1.9 | chaperone | 114 | 21-Dec-04 |
| 4avs | C5 | 1.4 | sugar binding protein | 204 | 19-Jun-13 |
| 4u62 | C5 | 1.55 | viral protein | 280 | 5-Aug-15 |
| 5a12 | C5 | 1.4 | oxidoreductase | 242 | 2-Sep-15 |
| 5lzh | C5 | 1.13 | toxin | 103 | 31-May-17 |
| 3h47* | C6 | 1.9 | viral protein | 231 | 23-Jun-09 |
| 4ox6* | C6 | 1.34 | structural protein | 127 | 27-Aug-14 |
| 2xf7 | C6 | 1.61 | viral protein | 51 | 11-Aug-10 |
| 1nlf | C6 | 1.95 | replication | 279 | 29-Apr-03 |
| 4w64 | C6 | 1.55 | Hcp1 protein (Unknown function) | 171 | 1-Jul-15 |
| 4owk | C7 | 2.0 | toxin | 138 | 28-May-14 |
| 1h64 | C7 | 1.9 | Sm-like protein | 75 | 19-Dec-02 |
| 4f87 | C8 | 1.4 | antimicrobial protein; viral protein | 72 | 25-Jul-12 |
| 3b8o | C8 | 2.4 | biosynthetic protein | 265 | 22-Jan-08 |
| 3zqo | C9 | 1.68 | DNA binding protein | 72 | 28-Dec-11 |
| 3p9a | C9 | 1.75 | DNA binding protein | 162 | 9-May-12 |
| 1orr | D2 | 1.5 | isomerase | 347 | 26-Aug-03 |
| 1zjz | D2 | 1.1 | oxidoreductase | 251 | 21-Jun-05 |
| 2bv4 | D2 | 1.0 | lectin | 113 | 25-May-06 |
| 4oqc | D2 | 1.3 | oxygen binding | 302 | 24-Dec-14 |
| 2vqr | D2 | 1.42 | hydrolase | 543 | 30-Sep-08 |
| 3v4f | D3 | 1.39 | signaling protein | 166 | 8-Aug-12 |
| 2bhq | D3 | 1.4 | oxidoreductase | 516 | 9-Mar-06 |
| 3qns | D3 | 1.4 | oxidoreductase | 353 | 27-Apr-11 |
| 2j5g | D3 | 1.46 | hydrolase | 263 | 16-Jan-07 |
| 1gxu | D3 | 1.27 | phosphatase | 91 | 12-Sep-02 |
| 2r8e | D4 | 1.4 | hydrolase | 188 | 23-Sep-08 |
| 3r1m | D4 | 1.5 | metal binding protein | 385 | 12-Oct-11 |

From the set of available structures, we randomly chose 10 cyclic complexes for testing flexible-backbone strategies. These complexes are listed with * after the PDB ID in the table above. The full benchmark could not be used as ensemble generation and global docking starting with 150+ monomer conformations were extremely resource consuming.

### Generation of homology-modeled monomers

For each protein, we obtained five homology-modeled monomers from the Robetta server (Song et al., 2013). We submitted the FASTA sequence for the first monomer chain to Robetta. To best simulate CAPRI conditions, we instructed Robetta to only consider templates older than the first REVDAT record for the complex being modeled, *i.e.* only those templates that would have existed before the PDB was deposited. Secondly, we instructed Robetta not to consider any symmetry information while modeling the monomers. The monomers obtained were used as input structures for the SymDock2 docking protocol.

### Generation of alternative conformations from the monomer

From the five monomer models produced for each target by Robetta, the model with the highest backbone RMSD less than 1.5 Å was selected as input for three alternative conformer generation methods in order to sample a variety of backbone conformations. This RMSD cutoff was chosen so that the conformations are not too close to the native structure and not so different that they do not fit in with other subunits. Relax, Backrub, and normal mode analysis (NMA) with perturbation steps of 1 Å were each used to produce 50 structures, resulting in an ensemble of 150 structures per target. The conformations in these ensembles were pooled with the homology-modeled monomers and used as input structures for the SymDock2 docking protocol.

### Relax

Rosetta Relax is a refinement protocol in which the protein is perturbed using a series of small backbone torsion moves (Tyka et al., 2011), which is followed by side-chain repacking and energy minimization along all torsion angles (*φ, ψ* and *χ* _i_). Each perturbation step is carried out at a particular weight of the van der Waals repulsive component of the all-atom score function (`fa_rep`). In each cycle, the weight is progressively ramped from 20% of the maximum (to allow atoms to come closer) to 100% (to resolve clashes). The lowest energy structure after 20 such cycles is chosen as the final structure for that trajectory. Relax was used to generate 50 monomer conformations.

The Relax protocol was implemented using the following command:

~~~
relax.linuxgccrelease
   -in:file:s <PDB> -nstruct 50 -relax:thorough
~~~

### Backrub

Rosetta Backrub attempts to capture small conformational changes that proteins undergo in solution (Smith and Kortemme, 2008). The protein backbone is divided into segments and each segment is rotated about the axis joining the first and the last backbone atom of the segment, while fixing the rest of the protein. The rotational movements are along six internal backbone degrees of freedom: the *φ, ψ*, and the N-C_α_-C bond angles at each pivot. This is followed by side-chain repacking and energy minimization along all torsion angles (*φ*, *ψ* and *χ* _i_). This process is repeated for 20,000 trials and the lowest energy structure is chosen. Backrub was used to generate 50 monomer conformations.

The Backrub protocol in Rosetta was implemented using the following command:

~~~
backrub.linuxgccrelease
   -in:file:s <PDB>-backrub:mc_kt 0.6
   -nstruct 50 -backrub:ntrials 20000
~~~

### Perturbation along normal modes

A normal mode of the protein is a collective motion in which all bonds are vibrating with the same frequency and phase: normal mode analysis (NMA) reveals accessible low-frequency vibrational modes that are thought to capture biologically relevant protein motions (Go et al., 1983). In Rosetta, NMA is implemented via the XML interface RosettaScripts (Fleishman et al., 2011). To generate monomer conformations, the protein is perturbed by steps of 1 Å randomly distributed over the first five normal modes. As this motion disrupts bond angles and bond lengths, the perturbed structure is subsequently relaxed using the aforementioned method with the exception of energy minimization being along Cartesian coordinates of the atoms. The score function of Relax is biased to favor ideal bond angles and bond lengths. This method was used to generate 50 monomer conformations.

The following command was used for running NMA-Relax in Rosetta:

~~~
rosettascripts.linuxgccrelease
     -in:file:s <PDB> -nstruct 50 -parser:protocol nma.xml
~~~

In this command, nma.xml contains the details of the protocol which is outlined below:

~~~
<ROSETTASCRIPTS>
   <SCOREFXNS>
       <ScoreFunction name="ref_cart" weights="ref2015_cart" />
   </SCOREFXNS>
   <RESIDUE_SELECTORS>
   </RESIDUE_SELECTORS>
   <TASKOPERATIONS>
   </TASKOPERATIONS>
<FILTERS>
   </FILTERS>
   <MOVERS>
       <NormalModeRelax name="nma" cartesian="true"
       centroid="false" scorefxn="ref_cart" nmodes="5"
       mix_modes="true" pertscale="1.0" randomselect="false"
       relaxmode="relax" nsample="20"
       cartesian_minimize="false"
   />
   </MOVERS>
   <APPLY_TO_POSE>
   </APPLY_TO_POSE>
   <PROTOCOLS>
      <Add mover="nma" />
   </PROTOCOLS>
   <OUTPUT scorefxn="ref_cart" />
</ROSETTASCRIPTS>
~~~

### Symmetry Definitions

We symmetrized the monomeric input structure using symmetry definitions in Rosetta’s symmetry framework (DiMaio et al., 2011). We used two kinds of symmetry definitions: general (also called *de novo*), and specific. Symmetry definitions contain information about the rigid-body arrangement of the subunits, how to scale the energy from calculations on one subunit (or a set of subunits), and specification of the degrees of freedom for the system.

We generated general or de novo symmetry definitions for each point symmetry in the benchmark using the pre-packaged Rosetta script, make_symmdef_file_denovo.py. This script inputs the symmetry group (Cn or Dn) along with the number of subunits. No information about any specific PDB is supplied. We used these definitions for global docking simulations. For example, the following command generates a general C2 symmetry definition:

~~~
make_symmdef_file_denovo.py –symm_type cn –nsub 2 > C2.symm
~~~

We generated specific symmetry definitions for each complex using the pre-packaged Rosetta script, make_symmdef_file.pl. This script inputs a symmetric PDB along with chain ID’s of the principal subunit (A), an interacting subunit (B), and in case of dihedrals, the chain ID of a chain in the sub-system in which A is not present (X). We also specified that we were using non-crystallographic symmetry (NCS) and the farthest interacting subunits were at a distance of d Å. We used these definitions for local docking, bound re-docking and bound refinement. The following command generates a specific symmetry definition from <PDB>:

~~~
make_symmdef_file.pl –m NCS –p <PDB> –a A -i B X –r d -f > <PDB>.symm
~~~

The symmetry definitions were used as inputs for the SymDock and SymDock2 protocols.

### Global docking simulations

We performed global docking using general symmetry definitions for the point symmetry of the given complex. The five homology-modeled monomers were used as inputs, each of which was used to start 1,000 independent trajectories to generate a total of 5,000 models. To evaluate backbone flexibility using conformational selection, the five monomer conformations were supplemented with another 150 conformations generated using Relax, Backrub and NMA. For these simulations, each input conformation was used to start 500 trajectories, thus generating a total of 77,500 models.

Global docking simulations with SymDock/SymDock2 used the following command:

~~~
SymDock.linuxgccrelease
   -in:file:l <list_of_input_PDBs>-nstruct (where n = 500 or 1000)
   -symmetry:symmetry_definition <general_symm_def_file>
   -symmetry:initialize_rigid_body_dofs
   -symmetry:symmetric_rmsd
   -ex1 -ex2aro -out:file:fullatom
~~~

To run just the coarse-grained phase, we removed

~~~
   -ex1 -ex2aro -out:file:fullatom
~~~

In SymDock2, the following options were added to enable Motif Dock Score:

~~~
   -docking_low_res_score motif_dock_score
   -mh:path:scores_BB_BB <Path to MDS tables>
   -mh:score:use_ss1 false-mh:score:use_ss2 false
   -mh:score:use_aa1 true-mh:score:use_aa2 true
~~~

### Local docking simulations

We performed local docking using specific symmetry definitions generated from the native PDB. The five homology-modeled monomers were used as inputs, each of which was used to start 1,000 independent trajectories. This generated a total of 5,000 models. The starting structure was randomly perturbed by 5 Å and 60° after symmetrization.

Local docking simulations with SymDock2 used the following command:

~~~
SymDock.linuxgccrelease
   -in:file:l <list_of_input_PDBs>-nstruct 500
   -symmetry:symmetry_definition <specific_symm_def_file>
   -symmetry:initialize_rigid_body_dofs
   -symmetry:symmetric_rmsd
   -symmetry:perturb_rigid_body_dofs 5 60
   -ex1 -ex2aro -out:file:fullatom
   -docking_low_res_score motif_dock_score
   -mh:path:scores_BB_BB <Path to MDS tables>
   -mh:score:use_ss1 false-mh:score:use_ss2 false
   -mh:score:use_aa1 true-mh:score:use_aa2 true
~~~

### Bound re-docking

In order to assess the ability of the coarse-grained phase in SymDock and SymDock2 to correctly identify and score near-native subunit arrangement, we re-docked the bound conformation. We started with the native monomer and specific symmetry definition taken from the native PDB, and ran the coarse-grained phase using the following command:

~~~
SymDock.linuxgccrelease
   -in:file:s <PDB_with_native_chain_A> -nstruct 100
   -symmetry:symmetry_definition <specific_symm_def_file>
   -symmetry:initialize_rigid_body_dofs
   -symmetry:symmetric_rmsd
~~~

In SymDock2, the following options were added to enable Motif Dock Score:

~~~
   -docking_low_res_score motif_dock_score
   -mh:path:scores_BB_BB <Path to MDS tables>
   -mh:score:use_ss1 false-mh:score:use_ss2 false
   -mh:score:use_aa1 true-mh:score:use_aa2 true
~~~

### Bound refinement

In order to assess the shape of the energy landscape near the bound conformation, we started with the native structure and perturbed it by iteratively re-packing side chains and minimizing energy along torsion angles at the interface and along the inter-subunit distance. We started with the native monomer and specific symmetry definition taken from the native PDB, and ran only the all-atom refinement protocol using the following command:

~~~
SymDock.linuxgccrelease
   -in:file:s <PDB_with_native_chain_A>-nstruct 100
   -symmetry:symmetry_definition <specific_symm_def_file>
   -symmetry:initialize_rigid_body_dofs
   -symmetry:symmetric_rmsd
   -docking:docking_local_refine
   -ex1 -ex2aro -out:file:fullatom
~~~

### Filtering Docking Models

Both Rosetta SymDock and SymDock2 filter out demonstrably poor models after the coarse-sgrained stage and after refinement (see Figures S1 and S2). The low-resolution filter after the coarse-grained phase in SymDock is based on terms of the centroid score function. This filter is `interchain_vdw` (penalizes clashes across chains) ≤ 1, `interchain_contact` (penalizes small interfaces) ≤ 10, and `atom_pair_constraint` (penalizes deviations from constraints) ≤ 1. For SymDock2, the low-resolution filter is based on MDS and is interchain_vdw ≤ 5. The high-resolution filter after all-atom refinement is more general and is common to SymDock and SymDock2. It is `total_score` ≤ 1,000,000 (total score of the model should not be ridiculously high) and `I_sc` ≤ 0 (it should be more favorable for the monomers to interact than remain separate).

### Simulation of Conformational Selection and Induced Fit

Once the ensembles are generated, in heterodimers, by superimposing different backbone conformations of a partner onto the current backbone along the interface, RosettaDock simultaneously samples rigid body orientations and backbone conformations (Marze et al., 2018). In SymDock, owing to multiple independent interfaces in homomers, this simultaneous sampling is not feasible since a conformation aligned with one interface may have significant clashes with the other interfaces. Thus, instead of sampling backbones during docking, we ran independent fixed-backbone simulations with each of the 155 monomer backbones. By creating 500 docked models per backbone, we generated a total of 77,500 models per complex. As the total number of docked models was different when using just the homology-modeled monomers and when supplementing it with the ensemble, we simulated the selection of 2,500 models for analysis. (If we had only generated 500 models starting from the five homology-modeled monomers, we would have obtained 2,500 docked models; hence, this value.) Unlike conformational selection, simulating induced fit does not require a multitude of backbone conformations to be sampled independently. Starting with just the five homology-modeled monomers, we generated 5,000 models. For an even comparison, we simulated the selection of 2,500 models for analysis. Thus, not only does inducing a fit after the coarse-grained phase improve the docking performance, it requires far fewer models than conformational selection to capture relevant backbone motion.

## Quantification and Statistical Analysis

### Binding energy funnel characterization

We compared the characteristics of the binding energy funnel in the 43 homomeric complexes in this study with 87 hetero-dimers previously studied (Marze et al., 2018). After fixed-backbone refinement of the native structure, the slope of the binding funnel is defined as the slope of the least-squares fit line for all models under 2 Å RM_Cα_ from the native complex. For homomeric complexes, 21 of the 43 complexes examined converged to the same state for all models (not necessarily at zero RMSD_Cα_), but none of the 87 hetero-dimeric complexes did so, which demonstrated the narrowness of the funnel in homomeric complexes. For 16 of the remaining homomeric complexes and 60 hetero-dimeric complexes, where a binding funnel was recovered, the average values were calculated. As homomers generally have extensive interfaces owing to multivalent interactions, we needed to normalize the values. Dividing by the number of subunits would not account for the fact that each subunit in a homomer has more interfaces than a heterodimer. The number of interfaces is difficult to define as different homomers have different extents of interactions with non-neighboring interfaces. Instead, we normalized the slopes by dividing them by the lowest interface score observed for the complex and compared the normalized values. The radius of the funnel is defined as the difference between the models with the largest and the smallest interface RMSD_Cα_ from the native structure. For the complexes in which all models converged to the same RMSD_Cα_, the funnel radius is zero. While calculating average funnel radius, we excluded complexes for which funnels could not be recovered, but included complexes with a zero funnel radius.

### Bootstrapping

As SymDock and SymDock2 rely on random moves to dock homomers, the final output model of each trajectory is different. To produce more information about the underlying distribution of each success metric, we resample with replacement from the available model set. Bootstrapping also allows us to compare results of runs where different number of models were generated. For example, when using conformational selection from an ensemble of 155 conformations, we generated 77,500 models, but using induced fit refinement, we generated only 5,000 models. Resampling allows us to simulate selection of a desired number of models, which in this case was 2,500 models. For the various success metrics, we reported medians, averages, and standard deviations across 1,000 re-sampling attempts.

### Success evaluation criteria

To evaluate the success of the docking simulations on the symmetric benchmark targets, we used two kinds of metrics: a near-native model count in the top-scoring models (*N#*) and fold-enrichment of near-native models in the low-scoring set (*E*_*N%*_). For example, *N5*, *N50* and *N500* are the number of near-native models in the 5, 50 and 500 top-scoring models, respectively. Fold-enrichment in the N% top-scoring models is defined as:

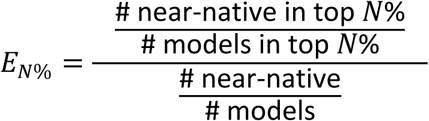

The bootstrapped averages for the success metrics are denoted by ⟨·⟩.

For coarse-grained models, near-native is refined as RMSD_Cα_ ≤ 5 Å. For all-atom models, near-native is defined as acceptable, medium-quality, or high-quality as per the CAPRI criteria listed below, which are based on the ligand RMSD_bb_, interface RMSD_bb_, and fraction of native contacts recovered (Méndez et al., 2003).

| Rank | Criteria |
| --- | --- |
| High-quality | $F_{\text{nat}} \geq 0.5$ and ($L_{\text{rmsd}} \leq 1.0$ or $I_{\text{rmsd}} \leq 1.0$) |
| Medium-quality | $F_{\text{nat}} \geq 0.3$ and ($1.0 < L_{\text{rmsd}} \leq 5.0$ or $1.0 < I_{\text{rmsd}} \leq 2.0$) |
| Acceptable | $F_{\text{nat}} \geq 0.1$ and ($5.0 < L_{\text{rmsd}} \leq 10.0$ or $2.0 < I_{\text{rmsd}} \leq 4.0$) |
| Incorrect | $F_{\text{nat}} < 0.1$ or ($L_{\text{rmsd}} > 10.0$ and $I_{\text{rmsd}} > 4.0$) |

After the full protocol, if ⟨*N5*⟩ ≥ 3, *i.e.* if, on average, at least 3 of the 5 top-scoring models are near-native, the complex is said to be successfully docked. This criterion was relaxed to ⟨*N10*⟩ ≥ 1 when comparing to other methods to allow a fair comparison.

### Data and Software Availability

As a part of the Rosetta software suite, Rosetta SymDock2 is available at https://www.rosettacommons.org to all non-commercial users for free and to commercial users for a fee.

